# Altered small-world functional network topology in patients with optic neuritis: A resting- state fMRI study

**DOI:** 10.1101/2020.06.09.141432

**Authors:** Ke Song, Juan Li, Yuanqiang Zhu, Fang Ren, Lingcan Cao, Yi Shao, Zi-Gang Huang

## Abstract

**Purpose:** This study investigated changes in small-world topology and brain functional connectivity in patients with optic neuritis (ON) by resting-state functional magnetic resonance imaging (rs-fMRI) and based on graph theory.

**Methods:** A total of 21 patients with ON (8 males and 13 females) and 21 matched healthy control subjects (8 males and 13 females) were enrolled at the First Affiliated Hospital of Nanchang University and underwent rs-fMRI. Data were preprocessed and the brain was divided into 116 regions of interest. Small-world network parameters and area under the integral curve (AUC) were calculated from pairwise brain interval correlation coefficients. Differences in brain network parameter AUCs between the 2 groups were evaluated with the independent sample t-test, and changes in brain connection strength between ON patients and control subjects were assessed by network-based statistical analysis.

**Results:** In the sparsity range from 0.08 to 0.48, both groups exhibited small-world attributes.

Compared to the control group, global network efficiency, normalized clustering coefficient, and small-world value were higher whereas the clustering coefficient value was lower in ON patients. There were no differences in characteristic path length, local network efficiency, and normalized characteristic path length between groups. In addition, ON patients had lower brain functional connectivity strength among the rolandic operculum, medial superior frontal gyrus, insula, median cingulate and paracingulate gyri, amygdala, superior parietal gyrus, inferior parietal gyrus, supramarginal gyrus, angular gyrus, lenticular nucleus, pallidum, superior temporal gyrus, cerebellum_Crus1_L, and left cerebellum_Crus6_L compared to the control group (P < 0.05).

**Conclusion:** The brain network in ON has a small-world attributes but shows reduced and abnormal connectivity compared to normal subjects. These findings provide a further insight into the neural pathogenesis of ON and reveal specific fMRI findings that can serve as diagnostic and prognostic indices.

## Introduction

Optic neuritis (ON) is a condition affecting 115 out of every 100,000 persons^[1]^**Error! Reference source not found.** that is characterized by inflammation and demyelination of the optic nerve as a result of infection or systemic autoimmune disease. The main clinical symptoms are sudden loss of visual acuity in one or both eyes within a short period of time, relative afferent pupil disorder (RAPD), papillary edema, pain during eye rotation, and visual field defect^[2]^**Error! Reference source not found.**. ON is closely related to demyelinating diseases of the central nervous system such as optic neuromyelitis and multiple sclerosis, among others^[3]^. Thinning of the retinal nerve fiber layer around the optic papilla in ON is observable by optical coherence tomography. ON can cause severe visual impairment but the pathogenesis is not fully understood, although it is thought to involve inflammation or immune factors that lead to optic nerve damage and ganglion cell apoptosis. In addition to demyelination, ON patients have abnormal activity in many brain regions ^[4]^**Error! Reference source not found.**. For example, brain atrophy was observed in patients with chronic recurrent solitary ON^[5]^, along with Wallerian degeneration in the optic tract, cerebellum, thalamus, posterior cingulate, and other brain areas^[6]^, indicating that specific brain areas are affected in ON.Given that the visual loss caused by ON can negatively affect the quality of life of patients, it is important to clarify the pathogenesis and associated changes in the brain. Resting-state functional magnetic resonance imaging (rs-fMRI) is a safe and widely used method for evaluating brain activity^[7]^ based on detection of the balance between local separation and global integration of signals associated with interconnected neurons. The normal human brain network has a short path length and high transmission efficiency, known as small-world attributes^[8]^**Error! Reference source not found.**; these enable the brain network to meet local and global demands and balance functional integration and separation in order to achieve synchronization of neural activity between different brain regions. Thus, small-world attributes allow efficient information transmission at low wiring cost^[9]^.

Graph theory, which is the study of the topologic structure of networks, provides a means of quantifying white matter changes in the brain^[10]^by considering these as a set of general elements sharing a specific relationship beyond anatomic connections^[11]^**Error! Reference source not found.**. Graph theory can be applied to rs-fMRI to characterize the functional connectivity and obtain a structural map of the brain from functional data, which can provide insight into the anatomic basis of brain dysfunction^[12]^and thus serves as an important reference for the diagnosis and treatment of diseases. Graph theory analysis has been widely used in studies on the mechanisms of post-traumatic stress disorder^[13]^**Error! Reference source not found.**, Alzheimer’s disease^[14]^, schizophrenia^[15]^ **Error! Reference source not found.**, stroke^[16]^, epilepsy^[17]^, and concussion^[18]^.

Most previous studies on brain abnormalities in ON have focused on anormal activity in specific brain regions. However, changes in small-world attributes and brain functional connectivity caused by ON have not been assessed. This was investigated in the present study by comparing small-world attributes and brain network connectivity in patients with ON and normal subjects by rs-MRI and the application of graph theory.

## Materials and Methods

### Subjects

A total of 21 ON patients (8 males and 13 females) were recruited at the Department of Ophthalmology, the First Affiliated Hospital of Nanchang University Hospital. Inclusion criteria were as follows: 1) sudden loss of visual acuity in one or both eyes within a short period of time; 2) positive RAPD or abnormal visual evoked potentials; 3) no visual field abnormalities related to nerve fiber injury; 4) None of the subjects had a history of psychiatric or neurological disorders; 5) no acute visual loss caused by other ophthalmologic or nervous system diseases; 6) no history of mental disorders, diabetes, hypertension, taking psychotropic drugs; 7) no history of drug, smoking, or alcohol addiction; and 8) average somatotype and weight. (**Figure 1**) We also recruited 21 age-, sex-, and weight-matched healthy control (HC) subjects (8 males and 13 females) who met the following criteria: 1) no pathway or brain parenchyma abnormalities observed by head MRI; 2) no ophthalmic disease and maximum corrected visual acuity >1.0; 3) no neuropsychiatric abnormalities or headache; and 4) no contraindications for MRI. After being informed of the nature of the study, all patients (or their guardians for participants under 18 years old) provided written, informed consent before participating. The study was approved by the research ethics committee of the First Affiliated Hospital of Nanchang University and the protocol was in accordance with the Helsinki Declaration.

**Figure 1.**
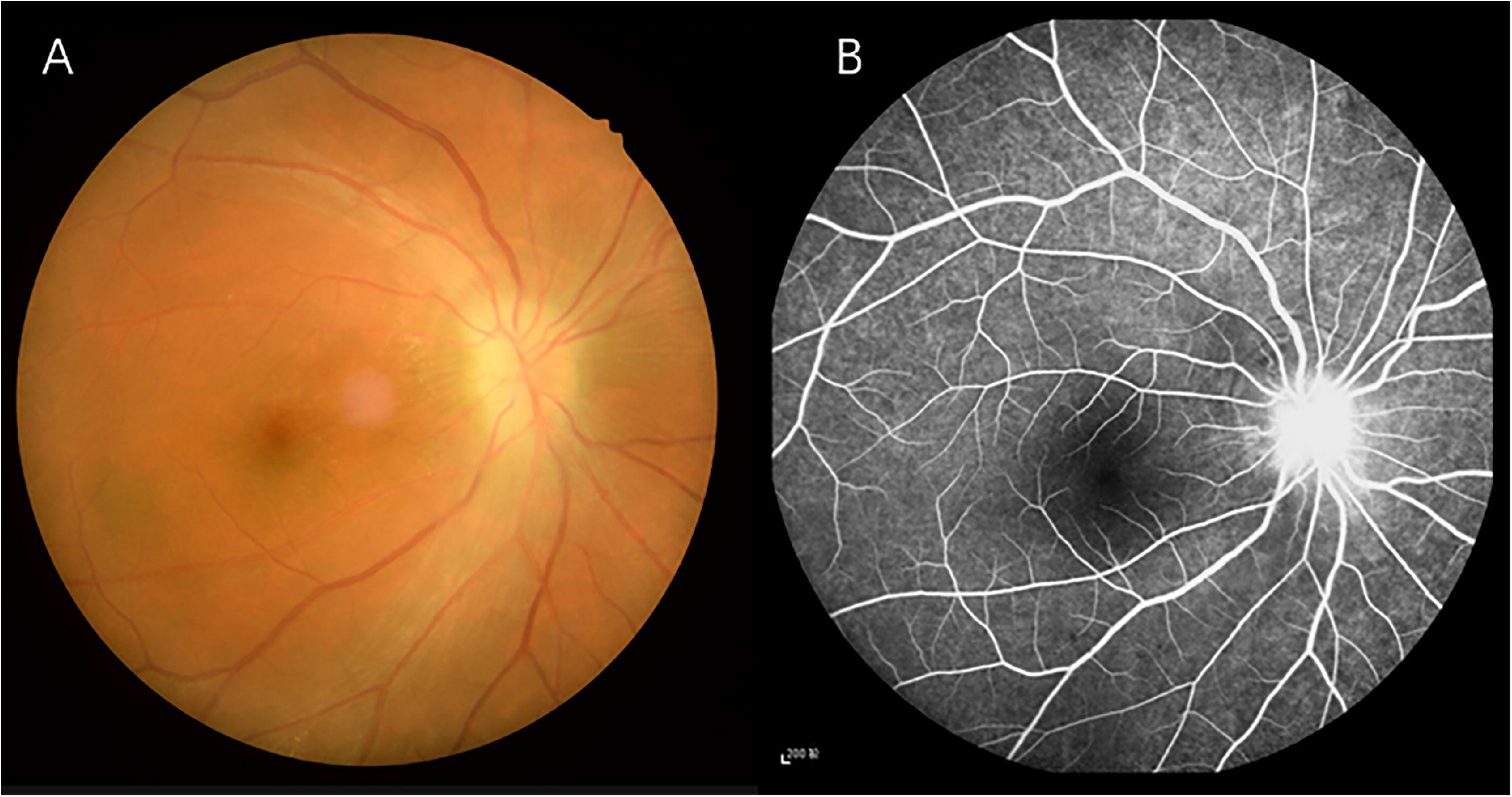
Eye examination data of ON patients. **Notes:**Figure A shows the results of fundus photography in ON patients. Figure B shows the results of fundus fluorescein angiograp(FFA) in ON patients. **Abbreviations:** ON, optic neuritis; FFA, fundus fluorescein angiograp.

### MRI data acquisition

A 3.0T TrioTim (Siemens, Munich, Germany) MR scanner and 8-channel head coil were used to collect rs-fMRI data and 3-dimensional high-resolution T1-weighted anatomic images. Participants were instructed to avoid drinking alcohol or coffee before the scan and those with intracranial lesions were excluded. During the scan, the subjects lay quietly with their eyes closed, breathe evenly and remain still, and avoid any mental activities insofar as possible. The subject was used a sponge pad to fix the head to reduce head movement, and the subject wore earplugs to block noise. For rs-fMRI, the parameters were as follows: repetition time (TR) = 2000 ms, echo time (TE) = 40 ms, flip angle = 90°, field of view (FOV) = 230mm × 230mm, matrix = 64 × 64, slice thickness = 4 mm, and slice number = 240 mm.The scanning parameters for T1-weighted structural images were as follows: TR = 1900 ms, TE= 2.26 ms, flip angle = 9°,FOV = 250 × 250mm, matrix = 256 × 256, layer thickness = 1 mm, layer number = 176.

### MRI data preprocessing

In order to eliminate the impact of magnetic field uniformity on network computing and intrasubject variability, poor-quality data were removed along with the first 10 time points. This study is based on MATLAB2014a (Mathworks,Natick,MA,USA) platform and uses DPARSFV2.3 software package to preprocess the data, The preprocessing involved in medicine standard, slice timing, time layer correction, spatial standardization, and smoothing with a 6×6×6mm full-width (half-height) Gaussian kernel^[19]^**Error! Reference source not found.**. Subjects with maximum x, y, or z displacement >1.5 mm and angular motion >1.5° were excluded from the analysis. Linear regression was used to remove nuisance variables containing signals from the region of interest (ROI) in the ventricles and areas centered on white matter. After correcting for head motion, the standard echo plane imaging template was used to normalize the fMRI image space to the Montreal Neurological Institute (MNI) space with resampling at a resolution of 3×3×3 mm. Finally, the data were then detrended to remove linear drift and temporally filtered by band-pass (0.01–0.08 Hz) to reduce the effects of low-frequency drift and high-frequency noise.

### Construction of brain network structure and analysis of topologic attributes

In order to obtain a whole-brain functional connectivity matrix for each subject, we divided the whole brain into 116 network nodes according to the anatomic automatic labeling template AAL116 (58 for each hemisphere, including the cerebellum). To define the ROI for the node region, the average processing time was calculated as the average of the fMRI time series of all voxels in the region, and the average time series of each region was obtained. The Pearson correlation coefficient between all possible pairs in the regional time series was calculated, obtaining a 116×116 correlation matrix each subject. We used a weighted matrix and included both positive and negative connections to construct a full-connection weighted network of the brain with sparsity as the threshold. The network analysis was carried out under a sparsity of 0.08–0.48 with an interval of 0.01. For the graph of each subject, we evaluated the whole-brain static network using this predefined range as the threshold with the following parameters: node clustering coefficient (Cp), characteristic path length (Lp), normalized clustering coefficient (γ), normalized characteristic path length (λ), small-world coefficient (σ) and brain network efficiency (the global efficiency (Eg), and the local efficiency (El)) ^[20]^. Network-Based Statistics (NBS) and GRETNA v2.0 (a toolbox for topological analysis of imaging connectomics) software were used to analyze network construction and assess differences in connectivity between the groups.

### Network-based statistical analysis

The brain is a complex network of functionally interconnected nodes that are distributed in a specific ROI^[21]^ Graph theory analysis was used to describe the topologic properties of networks but as it involved a large number of multiple comparisons, NBS provided by the false discovery rate (FDR) was used for whole-brain functional connectivity analysis at the ROI level and applied to connected components (subnets) that showed obvious differences between groups^[22]^.It was independently corrected for each connection in the network, and the corresponding P value for each link was independently calculated according to the strength of the paired association.

### Parameter integration

To evaluate overall differences between groups, the small-world parameters under each sparsity degree were integrated and the area under the curve (AUC) representing the overall level was recorded as aCp, aLp, aEg, aEl, aσ, aγ, and aλ

### Statistical analysis

The independent sample t-test (P<0.05 represented statistically significant differences) was used to evaluate differences in demographic and clinical variables between the ON and HC groups using SPSS v22.0 software (SPSS Inc, Chicago, IL, USA).Under a sparsity of 0.08–0.48, the independent sample t-test was used to assess small-world topologic differences in network metrics (aCp, aLp, aEg, aEl, aσ, aγ, and aλ), P<0.05 represented statistically significant differences. All normally distributed data are expressed as mean±standard deviation. We used NBS and link-based family-wise error rate (FWE) control provided by FDR to analyze potential brain functional connection differences and detect a contrast that was simulated between two groups, using independent a independent two-sample t test with P<0.05 and permutations of 5,000.

## Results

### Demographic and clinical characteristics

There were no statistically significant differences between the ON and HC groups in terms of weight (*P*=0.652), age (*P*=0.821), height (*P*=0.634), and BMI (*P*=0.963), while as the significant differences in VA-Right (*p*<0.001) and best-corrected VA-Left (*p*<0.001) between the two groups. (**Table 1**).

**Table 1.** Clinical characteristics of patients between ON and HC groups.

| characteristics | ON | HCS | *t*-value | *p*-values |
| --- | --- | --- | --- | --- |
| Male/female | 8\13 | 8\13 | NA | NA |
| Age(years) | 44.83±10.71 | 45.83±11.38 | -0.222 | 0.821 |
| Weight(kg) | 57.08±7.30 | 58.85±5.85 | -0.463 | 0.652 |
| Height(cm) | 160.81±9.31 | 161.38±6.28 | -0.485 | 0.634 |
| BMI(kg/m<sup>2</sup>) | 21.13±1.62 | 21.17±1.27 | -0.056 | 0.963 |
| Duration of ON(days) | 4.67±3.26 | NA | NA | NA |
| Duration from onset of ON to rs-fMRI scan(days) | 5.42±2.94 | NA | NA | NA |
| Best-correted VA,right | 0.25±0.32* | 1.30±0.31 | -8.138 | <0.001 |
| Best-correted VA,left | 0.85±0.52* | 1.28±0.32 | -2.481 | <0.001 |

**Notes:** Independent t-tests comparing the two groups (\* $P<0.05$ )represented statistically significant differences)
**Abbreviation:**ON,optic neuritis ;HCs,healthy controls;NA , not applicable;BMI,body mass index;rs-fMRI,resting-state functional magnetic resonance;VA,visual acuity.

### Analysis of small-world properties

Seven topologic small-world parameters were determined under a sparsity of 0.08 to 0.48 with an interval of 0.01. Cp, El, and Eg were positively correlated whereas Lp, γ, λ, and σ were negatively correlated with sparsity. Both groups had small-world attributes (λ>1, γ>1, σ>1). For the small-world indices, aγ(**Figure2B**) and aσ(**Figure2C**) were significantly higher for ON patients than for HC subjects (P<0.05). There were no statistically significant differences in aλ (**Figure2A**) between groups (**Table 2)**. For the other indicators of brain network topology, aCp (**Figure3A**) was lower whereas aEg (**Figure3B**) was higher in ON patients compared to HC subjects (both P<0.05). There was no significant difference in aEI(**Figure3C**) and aLp (**Figure3D**)between groups.(**Table 2**)

**Table 2.**
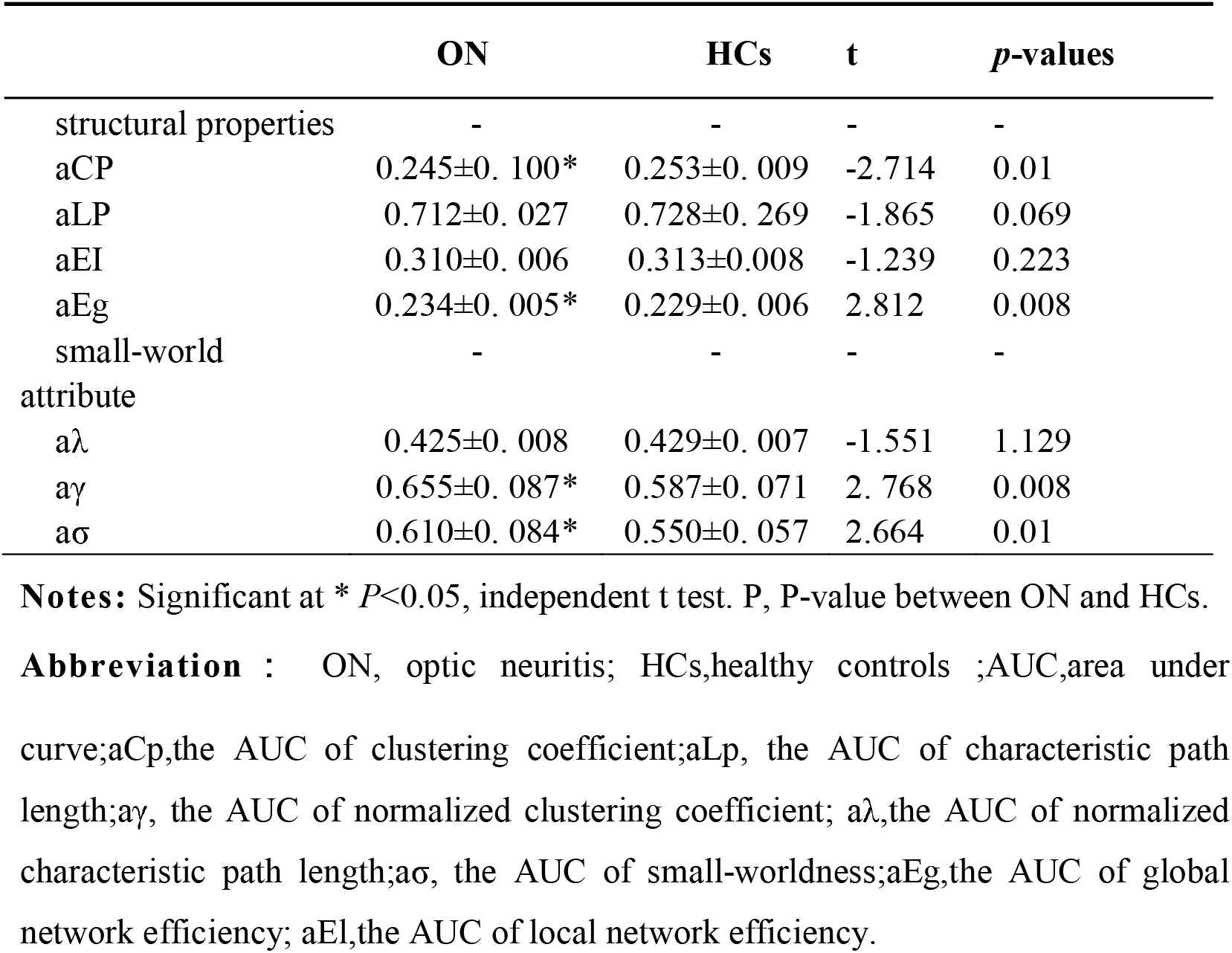
The AUC of the small-world parameters in patients with ONs and HCs.

**Figure 2.**
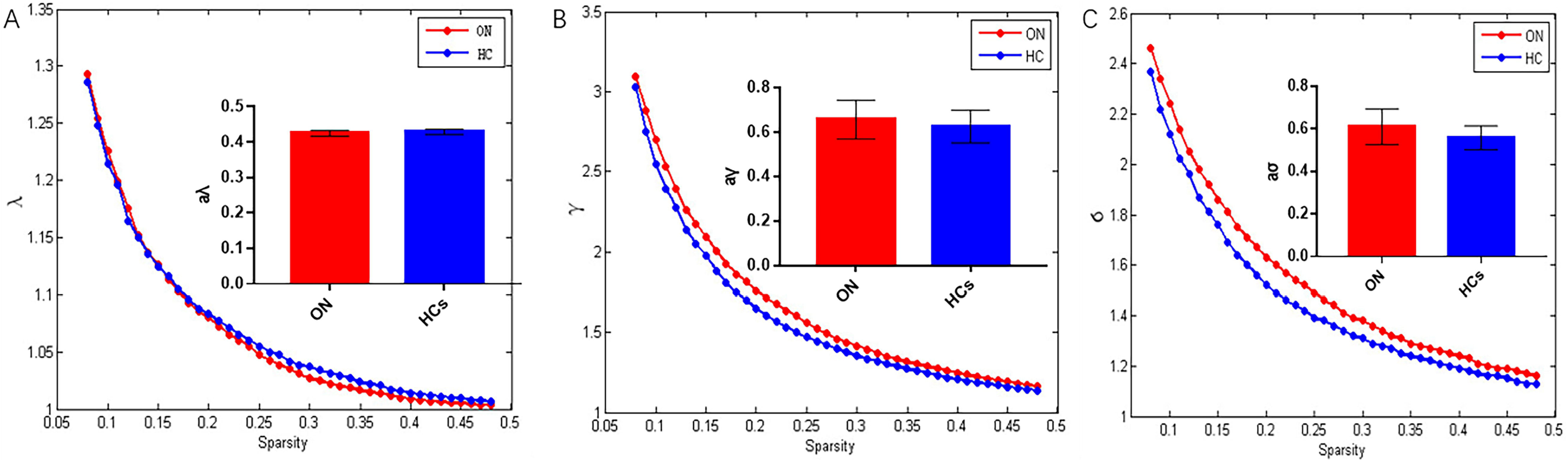
Comparison of analysis of small-world attribute of functional brain network between ON patients and HCs. **Notes:** Resting-state small-world parameter analyses showing that both ONs and HCs were consistent with small-world characteristics(λ>1, γ>1, σ>1). However, compared with the control group, the values of γ(B) and σ (C)in ON group increased significantly, and the difference was statistically significant (P<0.05). There was no significant difference in λ value (A) between the two groups (p>0.05). **Abbreviation**: ON, optic neuritis; HCs, healthy controls; AUC, area under curve; aγ, the AUC of normalized characteristic path length; aλ, the AUC of normalized characteristic path length; aσ, the AUC of small-worldness.

**Figure 3.**
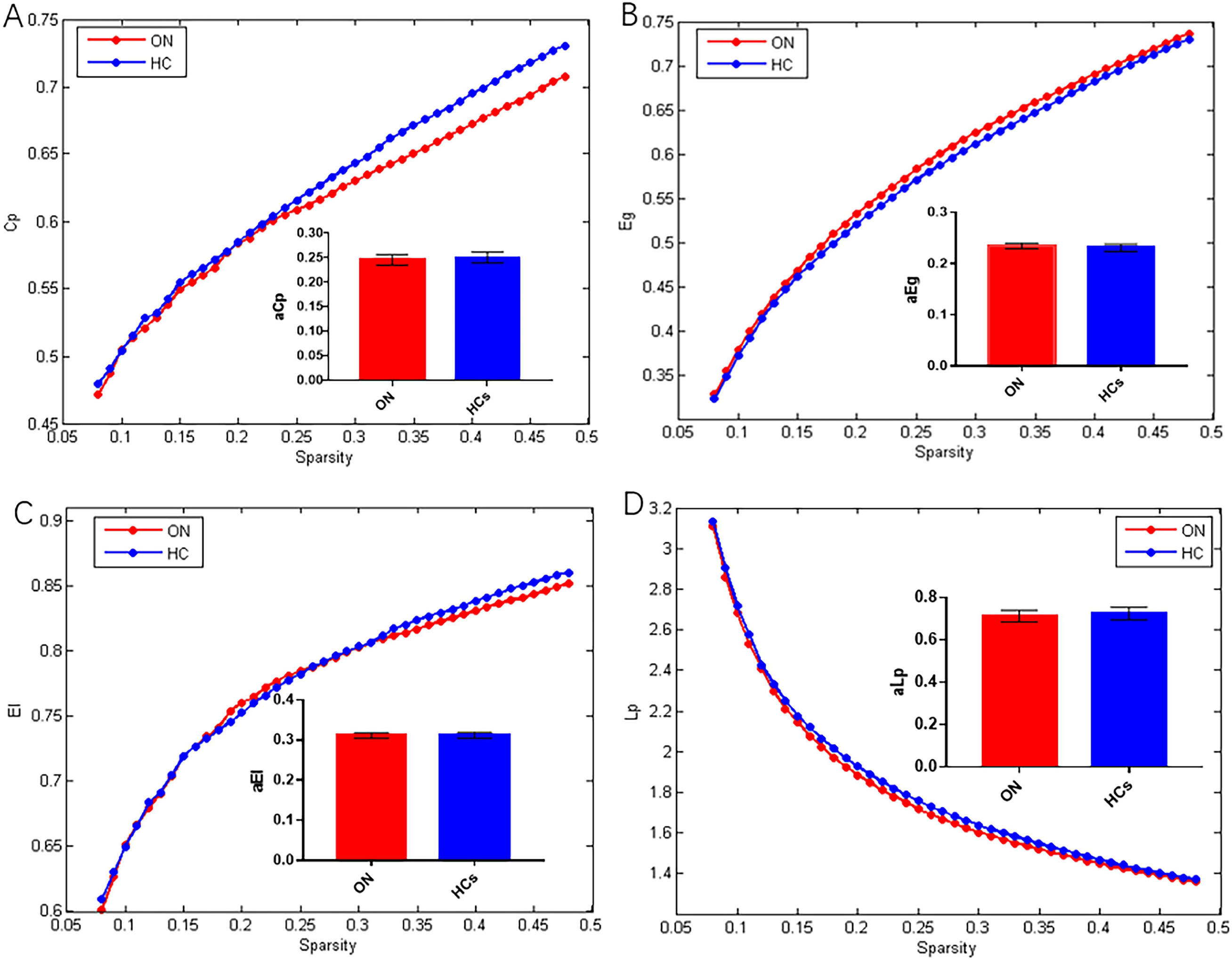
Comparison of structural properties of brain network between ON patients and HCs. **Notes:**Compared with HCs, the value of Cp in patients with ON was significantly lower (A) and the value of Eg was significantly higher (B), and the difference was statistically significant(P<0.05). There was no significant difference in the values of Lp (C)and EI (D) between the two groups (p>0.05). **Abbreviation**: ON, optic neuritis; HCs, healthy controls; AUC, area under curve; aCp, the AUC of node clustering coefficient;; aLp, the AUC of characteristic path length; aEg, the AUC of global network efficiency; aEl, the AUC of local network efficiency.

### Graph theory analysis of alterations in brain functional connectivity

Compared to the HC group, ON patients showed decreased brain functional connectivity. These mainly occurred among rolandic operculum(ROL), superior frontal gyrus, medial (SFGmed), insula(INS), median cingulate and paracingulate gyri (DCG), amygdala (AMYG), Superior parietal gyrus (SPG), inferior parietal gyrus, but supramarginal and angular gyri(IPL), supramarginal gyrus(SMG), angular gyrus(ANG), lenticular nucleus, pallidum (PAL), superior temporal gyrus(STG),cerebellum_Crus1(CRBLCrus1), cerebellum_Crus6 (CRBL 6). (**Figure4**) The difference is statistically significant (P<0.05). (**Table 3**)There were no instances where functional connectivity was higher in the ON group than in the HC group. These results demonstrate that optic nerve inflammation has a far-reaching effects on the functional brain network.

**Table 3.** Brain functional connectivity between ON patients and HCs identified by NBS analysis.

| **connectivity** | ***t-value*** | ***P<sub>NBS</sub>*** |
| --- | --- | --- |
| ROL.L to DCG.L | 6.026 | <0.05 |
| ROL.L to DCG.R | 5.747 | <0.05 |
| ROL.R to DCG.L | 5.239 | <0.05 |
| ROL.R to AMYG.R | 5.212 | <0.05 |
| SFG med.R to CRBL_Crus6_L | 5.205 | <0.05 |
| INS.L to DCG.L | 6.066 | <0.05 |
| INS.L to DCG.R | 5.169 | <0.05 |
| INS.L to AMYG.L | 5.284 | <0.05 |
| INS.L to SPG.L | 5.129 | <0.05 |
| INS.R to DCG.L | 5.454 | <0.05 |
| INS.R to IPL.R | 5.365 | <0.05 |
| INS.R to ANG.L | 5.071 | <0.05 |
| INS.R to ANG.R | 5.220 | <0.05 |
| INS.R to STG.L | 5.085 | <0.05 |
| SMG.L to PAL.R | 5.444 | <0.05 |
| SMG.R to CRBL_6_L | 5.257 | <0.05 |

**Notes:**NBS: $T > 3.92$ , $P < 0.05$ & 5,000 permutations , $P < 0.05$ indicates a significant difference between the groups.
**Abbreviation:** ON,optic neuritis ;HCs,healthy controls ; ROL, rolandic operculum; SFGmed , superior frontal gyrus,medial;INS, insula ; DCG , median cingulate and paracingulate gyri ; AMYG, amygdala; SPG , Superior parietal gyrus ; IPL , inferior parietal gyrus; SMG , supramarginal gyrus; ANG , angular gyrus ; PAL , lenticular nucleus,pallidum; STG , superior temporal gyrus ; CRBLCrus1 ,cerebellum\_Crus1; CRBL 6 , cerebellum\_Crus6; L, left hemisphere; R, Right hemisphere.

**Figure 4.**
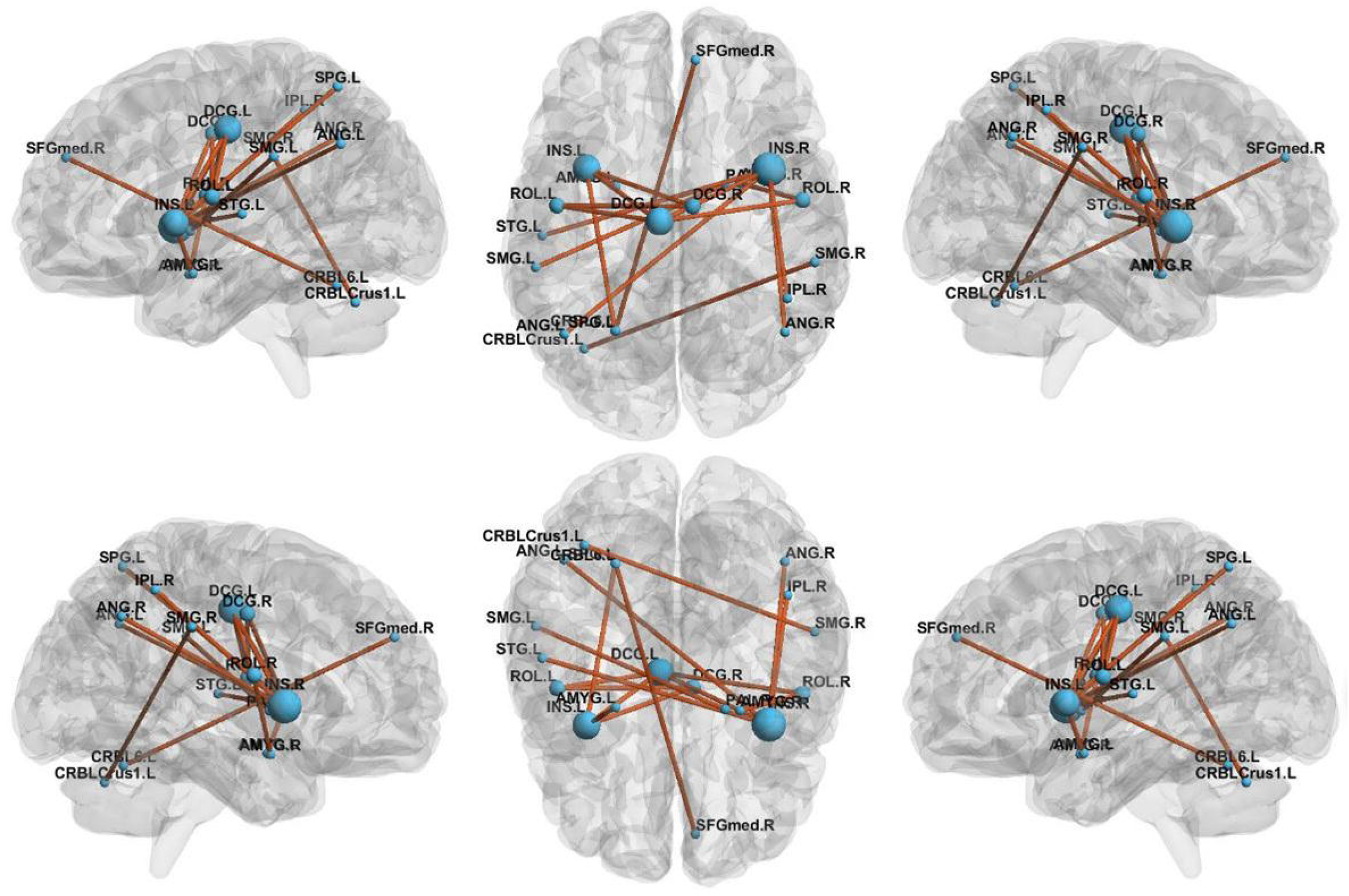
Graph theory analysis of alterations in brain functional connectivity. **Notes:**The figure shows the sub-network with decreased connectivity in ON patients compared to controls, identified by the NBS. Decreased brain functional connectivity in individuals with optic neuritis (ONs) compared to healthy controls. (NBS: T>3.92, P<0.05 & 5,000 permutations). **Abbreviation**: ON, optic neuritis; HCs, healthy controls; ROL, rolandic operculum; SFGmed, superior frontal gyrus, medial; INS, insula; DCG, median cingulate and paracingulate gyri; AMYG, amygdala; SPG, Superior parietal gyrus; IPL, inferior parietal gyrus, but supramarginal and angular gyri; SMG, supramarginal gyrus; ANG, angular gyrus; PAL, lenticular nucleus, pallidum; STG, superior temporal gyrus; CRBLCrus1, cerebellum_Crus1; CRBL 6, cerebellum_Crus6; L, left hemisphere; R, Right hemisphere.

## Discussion

This is the first study to use graph theory and NBS to analyze the small world and brain functional connectivity strength in ON. Inflammation and immune activation can cause demyelination of the optic nerve, leading to reduced signal transmission and visual impairment^[23]^**Error! Reference source not found.**. The demyelination and axon damage associated with ON was shown to result in the destruction of functional brain networks in a small-world study of craniocerebral trauma ^[24]^. In the present work, we found that ON patients retained small-world characteristics (λ>1, γ>1, and σ>1) although some network parameters were altered. Compared to HCs, Eg, γ, and σ were increased whereas Cp was decreased in ON patients; moreover, Cp, El, and Eg were positively correlated and Lp, γ, λ, and σ were negatively correlated with sparsity. The lower Cp in ON patients may reflect a reduced functional connectivity in some brain regions caused by extreme demyelination, which has also been observed in cases of axonal injury. Patients with long-term disturbance of consciousness also show alterations in small-world parameters. Lp measures the capacity for global information transmission and is related to cognitive function ^[9,25]^, while Eg represents global network efficiency. In our study, Eg was higher in ON patients than in HC subjects, suggesting greater efficiency in network information transmission.Patients with ON often have monocular disease^[26]^**Error! Reference source not found.**. Insufficient stimulation of the visual cortex from the decreased visual acuity in one eye can lead to compensatory activation of the contralateral brain region ^[27]^. We observed an increase in the amplitude of low-frequency fluctuation (ALFF) value of the left superior temporal gyrus in ON patients, consistent with findings from other MRI studies of ON^[28]^. This implies that the increase in Eg in ON patients is a mechanism to offset brain network dysfunction^[29]^**Error! Reference source not found.**. The decreased λ value was accompanied by a compensatory reduction in Lp. The injury caused by ON alters small-world attributes of brain connectivity networks. The parameter σ is a clustering coefficient that represents approximate shortest path length ^[30,31]^. A lower σ represents a greater tendency toward a random brain network; in concussion, these were shown to be more susceptible to pathologic insults than small-world networks^[18]^. In this study, σ was >1 in ON patients, indicating that they have small-world characteristics; however, the value was higher than that in HC subjects. The optic nerve is rich in macrophages and T cells^[32,33]^; thus, the increase in σ may be attributable to inflammation or damage. γ is the standardized clustering coefficient and is used to measure dispersion within the network, with a higher value indicating a higher degree of grouping.

The significant difference in the static state connection network between ON patients and HC subjects emphasizes the role of NBS in multivariate comparisons. We observed a decrease in the connection strength of multiple brain regions in ON patients. The INS is located in the deep part of the lateral sulcus at the boundary between the annular sulcus and frontal, temporal, and parietal lobes. Abnormal activation of the insular-interstitial area has been reported in ON patients^[34]^ **Error! Reference source not found.**; additionally, the latency of visual evoked potentials decreased with REHO signal in the INS, which could decrease the connection strength between this and other brain regions^[35]^**Error! Reference source not found.**.The cerebellum is located in the inferior part of the brain posterior to the medulla oblongata and pons. Cerebellar demyelination has been reported in ON^[36]^, which could explain the reduced connectivity between the cerebellum and other brain regions in patients. The STG is located in the temporal lobe between the lateral and superior temporal sulci, and plays a key role in sound processing. An fMRI study revealed that the ALFF signal was decreased in the superior temporal gyrus of ON patients^[27]^**Error! Reference source not found.**, which may be related to the severity of ON. The STG is also involved in visual searching, and decreased magnetic resonance related signals in this region have been observed in patients with retinal detachment^[37]^ **Error! Reference source not found.**. An impaired STG in patients with ON could result in decreased connectivity with surrounding brain regions. The AMYG is located in the dorsomedial part of the anterior temporal lobe, slightly anterior to the top of the hippocampus and inferior horn of the lateral ventricle. As part of the limbic system, the AMYG plays an important role in generating, identifying, and regulating emotion^[38]^, and it is among the key brain areas responsible for normal and pathologic stress responses. A decreased connection strength between the AMYG and other brain areas in ON patients may indicate a reduced ability to respond to pathologic events, leading to the destruction of brain network structure^[39]^. The SMG contributes to the maintenance of short-term auditory-language, motor, and visual-spatial memory sequence^[39]^; the reduced connectivity between the SMG and surrounding areas in ON patients suggests that the normal perception of visual space is disrupted. The ANG, which is the visual language (reading) center, is arched around the end of the supratemporal sulcus in the temporal lobe. The ANG integrates incoming sensory and cognitive information, responds to stimuli in memory and learning^[40]^, and functions in memory retrieval^[41]^. The activity of ANG-related neural circuits is increased during eye-to-eye communication, and both the ANG and STG have been implicated in Wernicke’s (sensory) aphasia ^[42, 43]^. Therefore, the decreased connectivity between the ANG and STG in ON may be associated with reading dysfunction. The SFG, located in the upper part of the prefrontal lobe, is involved in motor coordination, working memory, and resting-state and cognitive control. The fractional ALFF signal of the SFG was shown to be positively correlated with perceived stress^[44]^, and the gray matter structure of the SFG has been implicated in the processing of early and recent life stress events^[44,45]^. The reduced SFG connection strength in ON patients may be due to stress caused by optic nerve inflammation. The SPG is located in the dorsomedial parietal lobe anterior to the parietal-occipital sulcus and above the parietal sulcus. In the posterior part of the retrocentral sulcus, the SPG participates in stereoscopic visual processing^[46]^ and plays a key role in defining visual space in language and motor areas^[47]^**Error! Reference source not found.**. Additionally, the SPG controls eye movement^[48]^**Error! Reference source not found.**. Eye rotation pain is common in ON^[49]^, and may also be associated with changes in brain connectivity; however, in our study, the strength of the connection between the SPG and surrounding brain regions was decreased, suggesting that there was damage to the area corresponding to eye movement pain. The PAL is located in the lentiform nucleus of the striatum. Lesions involving the extrapyramidal system and pyramidal tract may cause movement disorder and nystagmus in the eyes. The DCG, which is crescent-shaped and surrounds the corpus callosum, is a major component of the limbic system^[50]^ and is related to memory and spatial orientation^[51]^**Error! Reference source not found.**. The reduced connection strength between the DCG and other brain areas in patients with ON could affect their capacity for spatial localization. ROL is the cortex adjacent to the insular, which is one of the major regions involved in the language processing system^[52]^,and it also involves in motor, sensory, autonomic and cognitive processing^[53]^Relevant research data also confirms the role of rolandic operculum and neighboring areas (such as insular) in processing sensory signals related to other conscious operations (such as visual awareness)^[54]^. The decrease of the connection strength between ROL and the surrounding brain area may reflect the impairment of visual function. IPL is a part of the parietal lobe and is related to visual recognition and selective scanning targets^[55]^**Error! Reference source not found.**. Finally, the observed changes in the IPL imply the impairment of stereoscopic visual function**Error! Reference source not found.**^[54]^. Taken together, these results demonstrate that optic nerve inflammation has far-reaching effects on the functional brain network.

### Limitations and strengths

There were some limitations to this study. Because of the small sample size, we did not examine the correlation between brain topologic characteristics and clinical manifestations of ON or between changes in brain structure and function. In the future, brain network changes in ON will be analyzed in a larger cohort by multimodal analysis.

## Conclusion

The results of this study show that the functional brain network of ON patients has small-world properties, but that these are significantly impaired relative to HC subjects. The changes in small-world properties observed in ON may be caused by demyelination resulting from inflammation and could reflect functional impairment in the brain. Our findings provide novel insight into the anatomic and functional brain abnormalities associated with ON, and also reveal specific fMRI findings that can serve as diagnostic and prognostic indices.

## Conflict of Interest Statement

This was not an industry supported study. The authors report no conflicts of interest in this work.

## Funding Disclosure

This research is supported by National Natural Science Foundation of China (No: 81660158, 81160118, 81400372, 81460092, 81500742); Medical Science Foundation of Guangdong Province (No:A2016184).Natural Science research Foundation of Guangdong Province (No:2017A030313614, 2017A020215187, 2018A030313117)

## Notes

### Competing Interest Statement

The authors have declared no competing interest.

